# Cell size homeostasis under the circadian regulation of cell division in cyanobacteria

**DOI:** 10.1101/2022.01.25.477786

**Authors:** Yuta Kitaguchi, Hajime Tei, Koichiro Uriu

## Abstract

Bacterial cells maintain their characteristic cell size over many generations. Several rodshaped bacteria, such as *Escherichia coli* and the cyanobacteria *Synechococcus elongatus*, divide after adding a constant length to their length at birth. Through this division control known as the adder mechanism, perturbation in cell length due to physiological fluctuation decays over generations at a rate of 2^−1^ per cell division. However, previous experiments have shown that the circadian clock in cyanobacteria reduces cell division frequency at a specific time of day under constant light. This circadian gating should modulate the division control by the adder mechanism, but its significance remains unknown. Here we address how the circadian gating affects cell length, doubling time, and cell length stability in cyanobacteria by using mathematical models. We show that a cell subject to circadian gating grows for a long time, and gives birth to elongated daughter cells. These elongated daughter cells grow faster than the previous generation, as elongation speed is proportional to cell length and divide in a short time before the next gating. Hence, the distributions of doubling time and cell length become bimodal, as observed in experimental data. Interestingly, the average doubling time over the population of cells is independent of gating because the extension of doubling time by gating is compensated by its reduction in the subsequent generation. On the other hand, average cell length is increased by gating, suggesting that the circadian clock controls cell length. We then show that the decay rate of perturbation in cell length depends on the ratio of delay in division by the gating *τ_G_* to the average doubling time *τ*_0_ as 2*^τ_G_/τ_0_^*^−1^. We estimated *τ_G_* ≈ 2.5, *τ*_0_ ≈ 13.6 hours, and *τ_G_*/*τ*_0_ ≈ 0.18 from experimental data, indicating that a long doubling time in cyanobacteria maintains the decay rate similar to that of the adder mechanism. Thus, our analysis suggests that the acquisition of the circadian clock during evolution did not impose a constraint on cell size homeostasis in cyanobacteria.

## 1 Introduction

A population of cells maintains a stable cell size distribution with well-defined mean and variance, due to the presence of cell size control mechanisms [1]. Cell size control is crucial especially for multi-cellular organisms, as their tissue size largely depends on cell size [2]. One of the most important determinants of cell size is division timing. Cell size control by division timing has been studied in prokaryotes [1, 3] and budding yeast [4, 5]. In rod-shaped bacteria such as *Escherichia coli* and *Bacillus subtilis*, the length on their long axis determines cell size [6]. In *E. coli* and other bacteria, constriction of a ring assembled with several proteins including FtsZ at midcell leads to cell division [7]. Recent experiments indicated that in many bacteria, doubling time and cell length at division, termed division length, differ between cells with different cell length at birth (birth length) [1, 6]. Rather, cells divide after adding a constant length to their birth lengths (Fig. 1A, B), and this division control is referred to as an adder mechanism [1, 3, 6, 8–10]. Previous imaging analysis of bacteria showed that the adder mechanism reduces cell length perturbation caused by transient physiological fluctuations at the rate 2^−1^ per generation (Fig. 1C; A) [6, 11]. Cell lengths of these bacteria, thus, are maintained near the species-specific mean length. In other words, the distribution of individual doubling times in these bacteria is unimodal [6].

**Figure 1:**
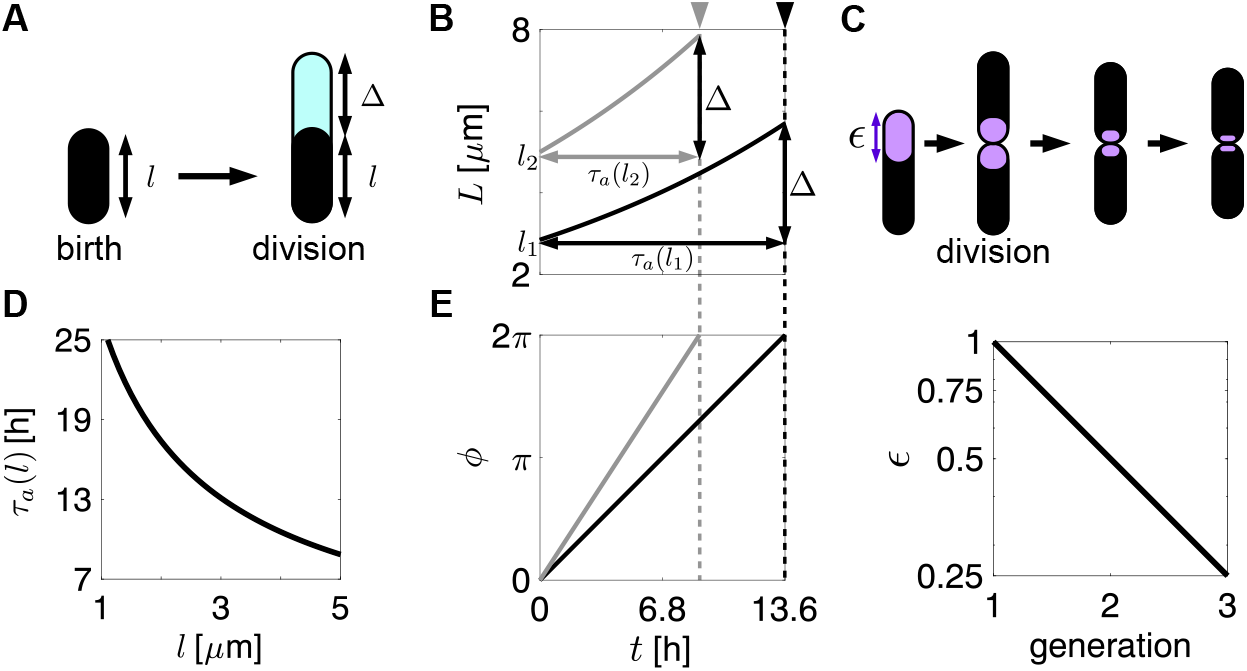
Adder mechanism. (A) A cell divides after the elongation of Δ regardless of its birth length *l*. (B) Cell length elongation from different birth lengths *l*_1_ (black) and *l*_2_ (gray). Cells add Δ to their birth length by the time when cell phase reaches *ϕ* = 2*π*. *τ_a_* is doubling time. (C) Decays of cell length perturbation *∊* (purple) over several generations. The cell length perturbation is decreased 2^−1^ per division with the adder mechanism. Vertical axis is a logarithmic scale. (D) Dependence of time *τ_a_* for a cell to add Δ as a function of its birth length *l*. (E) Dependence of cell phase progression on birth length. The black (gray) line indicates cell phase progression for a cell with a longer (shorter) birth length shown in (B). When *ϕ* = 2*π*, cells divide as indicated by arrowheads in (B). The cell phase is reset to 0 after division.

The all aforementioned bacteria divide regardless of environmental light conditions; their division frequency is uniform through the day-night cycle. In contrast, cell division in the photosynthetic cyanobacteria *Synechococcus elongatus* is strongly influenced by light conditions. Cyanobacterial cells cultured in a 12-hour light, 12-hour dark (12:12 LD) cycle elongate and divide only under light periods [12, 13]. To save their metabolic energy, these cells neither elongate nor divide under dark periods. Cyanobacteria’s strong dependence on light for cell division implies that predicting an appropriate time of day for division by an internal clock might be necessary for their survival [14].

Indeed, growing evidences show that cyanobacterial cell division is controlled by not only cell size and light signal, but also the circadian clock [12, 13, 15, 16]. The circadian clock in cyanobacteria consists of protein products of the *kaiA*, *kaiB* and *kaiC* genes. These three Kai proteins generate circadian rhythms in the phosphorylation state of KaiC proteins [17, 18]. KaiA binds to unphosphorylated KaiC and promotes its phosphorylation. After the dissociation of KaiA, KaiB associates with KaiC to promote its dephosphorylation. Along with the circadian phosphorylation cycle, the ATPase activity of KaiC also oscillates [19]. It peaks ~ 4 hours before the peak of KaiC phosphorylation at early subjective night [19]. Thus, the circadian clock consisting of the three Kai proteins determines the subjective time of cyanobacteria in constant light environment.

Previous experimental studies have shown that the cell division frequency in cyanobacteria when cultured under constant light (LL) decreases at early subjective night [12, 15]. In contrast, a clock mutant strain lacking *kaiBC* genes (Δ*kaiBC*) divides at a constant frequency throughout the day [13], suggesting that the circadian clock reduces cell division frequency at early subjective night in the wild type (Fig 2A). A reduction in cell division frequency at a specific time of the day has been termed circadian gating [12]. The detailed molecular mechanism underlying circadian gating is not yet known, but a previous experiment with various mutated KaiC proteins revealed that the ATPase activity of KaiC regulates cell division [16]. Specifically, KaiC with high ATPase activity activates the protein kinase RpaA, which then disrupts the localization of FtsZ protein to the midcell, resulting in the inhibition of cell division [16].

**Figure 2:**
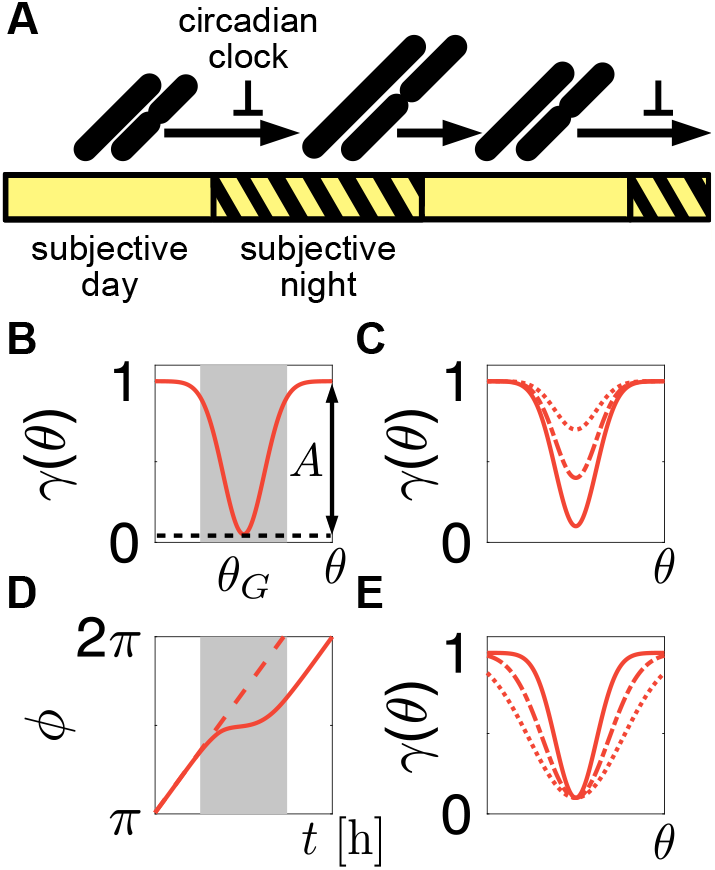
Circadian gating of cell division. (A) A circadian clock inhibits cell division at early subjective night under constant light (yellow). (B) Gating function *γ*(*θ*) in Eq. (5). (C) Gating function Eq. (5) with *A* = 0.3 (dotted), 0.6 (dashed) and 0.9 (solid). *β* = 50. (D) Deceleration of cell phase progression by the circadian clock via the gating function *γ*(*θ*) shown in (B) (solid red line). The dashed red line indicates the cell phase progression in the absence of the circadian regulation *γ*(*θ*) = 1 for comparison. (E) Gating function Eq. (5) with *β* =10 (dotted), 20 (dashed) and 50 (solid). *A* = 0.9.

Moreover, single-cell tracking of wild type cyanobacteria recently showed that a cell population in LL conditions can be classified into two subpopulations: one with a longer doubling time (16.3 hours on average) and the other with a shorter doubling time (11.6 hours) [13]. Consistent with the absence of circadian regulation of cell division, the Δ*kaiBC* mutant cells are classified into only a single group with an average doubling time of 13.6 hours [13]. Furthermore, quantitative analysis of cell length in the Δ*kaiBC* strain showed that the added length only weakly depends on birth length, suggesting that cell division in this mutant strain is determined by the adder mechanism [13]. Thus, the circadian clock in wild type cyanobacteria modulates cell division timing determined by the adder mechanism.

Based on these observations, previous theoretical studies inferred the effect of cyanobacterial circadian clock on cell division. Yang et al. 2010 fitted a mathematical model of the circadian gating to data on division timing and estimated the inhibition strength of cell division by the circadian clock [15]. They showed that the inhibition strength depends strongly on the phase of the circadian clock, but weakly on the phase of cell division cycle. In contrast, by using data on both cell length and doubling time, Martins et al. 2018 proposed a different model where the circadian clock promotes cell division at late subjective day, but inhibits division during the rest of day [13]. Recently, Ho et al. 2020 analyzed a model that describes dynamics of both cell size and the concentration of a divisor protein that after accumulating, leads to cell division [20]. In their model, the circadian clock modulates the production rate of the divisor protein, and thereby, changes the cell division frequency within a day.

These theoretical studies mainly focused on elucidating a functional form that describes the circadian regulation of cell division in a mathematical model. However, the dynamics of cell length and doubling time under circadian regulation remain unclear. For example, how does the combination of the adder mechanism and circadian gating generate cell populations with different doubling times as observed in recent experiments [13]? Furthermore, because cyanobacterial cells continue to elongate while cell division is inhibited by high KaiC ATPase activity [16], the circadian regulation of cell division may also enlarge the fluctuation in cell length. Hence, it is not obvious how cyanobacteria under LL conditions maintain cell size homeostasis.

To address these questions, we analyze cyanobacterial cell division when cultured under a LL condition using mathematical modeling and simulation. We incorporate the adder mechanism [21] into a previous model for circadian gating [15]. Simulations of this gating model show that an inhibition of cell division by the circadian clock at early subjective night increases added length and produces elongated daughter cells at late subjective night. These daughter cells grow faster due to their longer birth lengths and divide before the onset of the next gating. We show that the delay in cell division by gating increases both average birth length and added length, but, interestingly, does not affect average doubling time. Subsequently, we derive difference equations for birth length of cells in two successive generations. By performing a stability analysis on the difference equations, we show that the decay rate of perturbations to cell length is determined by the ratio of the delay in cell division caused by gating to the average individual doubling time. This ratio is close to zero under the LL condition, explaining the stable maintenance of cell length distribution. Taken together, the cyanobacterial circadian clock may be beneficial for energy control [14], but it can also decrease cell length stability by gating cell division. Our analysis reveals that the long average doubling time makes up for the adverse effects of the delay in cell division caused by circadian gating.

## 2 Model

Throughout this study, we consider a constant light condition where photosynthetic cyanobacteria can grow without interruption by a dark period. To analyze the effects of the circadian clock on cell length and doubling time, we develop a phenomenological model for the circadian regulation of cell division. Based on previous studies [12, 15], we assume that the circadian clock inhibits cell division at a certain time of day. We refer to this as the circadian gating of cell division. In addition, we include the adder mechanism where a cell divides after a constant length of elongation as observed in *E. coli*, *B. subtilis* [1, 6], and in cyanobacteria [10, 13]. The adder mechanism is independent of the circadian clock because it remains functional in the Δ*kaiBC* strain [13].

The model describes the time evolution of cell length *L* and cell phase *ϕ* that determines division timing. The elongation of cell length L is described as:

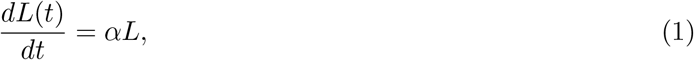

where *α* is the constant elongation rate (Fig. 1B). A previous experiment indicated that the elongation rate α of wild type cyanobacteria changes throughout the day even under constant light [13]. However, because the observed variation of the elongation rate was smaller compared to its time average [13], we assume that α is constant over time in the model for simplicity. After cell division, the cell length becomes *L_d_*/2 + *ξ* where *L_d_* is the cell length at division and *ξ* is a small random noise. In this study, we randomly choose the value of *ξ* from a uniform distribution between −0.01 and 0.01 *μ*m.

To determine division timing, we consider a cell phase variable *ϕ*. Physiologically, this cell phase *ϕ* is an abstract representation of the state of progress in DNA replication and accumulation of FtsZ for the next cell division. Accordingly, we set *ϕ* = 0 immediately after a cell division. Then, *ϕ* increases over time and the cell divides when *ϕ* = 2*π* (Fig. 1B, E). We describe the time evolution of *ϕ* between two successive divisions as:

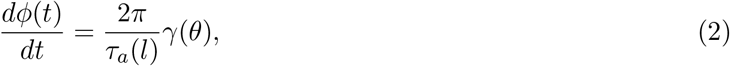

where *l* is the cell length at birth, termed birth length, *τ_a_*(*l*) is the doubling time determined by adder mechanism alone, *θ* is the phase of the circadian clock and *γ*(*θ*) is the modulation of speed of cell phase progression by the circadian clock described below.

In this study, we assume the adder mechanism for the cell length control. In the adder mechanism, a cell divides after adding Δ to its birth length l by elongation. Then, *τ_a_*(*l*) in Eq. (2) can be written as:

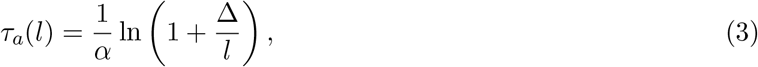

see A for derivation and ref. [21]. *τ_a_* becomes short as birth length *l* is large (Fig. 1D).

We model the influence of the circadian clock on cell phase *γ*(*θ*) based on a previous work (Fig. 2A) [15]. We assume that the phase of the circadian clock *θ* ∈ [0, 2*π*) increases at a constant rate *ω* = 2*π*/24:

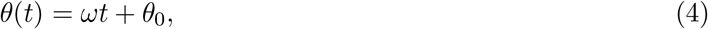

where *θ*_0_ is the initial phase of the circadian clock. We assign subjective time (circadian time, CT) for cells based on *θ*. Namely, *θ* = 0 is the onset of subjective day (CT = 0), and *θ* = *π* is the onset of subjective night (CT = 12). Then, we describe the modulation of cell phase progression by the circadian clock *γ*(*θ*) in Eq. (2) as:

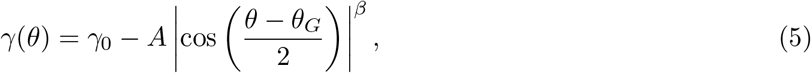

where *γ*_0_ is the basal regulation, A is the inhibition strength (A > 0), *θ_G_* is the gating phase denoting when the influence of the clock becomes strongest, and *β* is the exponent regulating gate width (Fig. 2B–E). A positive value of A results in deceleration of the cell phase progression at around *θ_G_* (Fig. 2C, D). Decrease in *β* widens the gate width (Fig. 2E). We refer to Eq. (5) as the gating function. Note that the influence of the circadian clock on cell phase is independent of the value of the cell phase itself. This simplification was validated by a previous quantification of the influence of the circadian clock on cell division [15].

Recent studies [13, 20] have suggested that circadian regulation of cell division might be more complex than gating [12]. These studies proposed that the circadian clock both promotes and inhibits cell division depending on the time of day. In this current study, however, we only consider the inhibition of cell division at a certain circadian phase for simplicity. We will validate this simplification in the results section by showing that the current model reproduces most of the statistics observed in the experimental data for cyanobacterial cell division. In addition, we argue that the inhibition of cell division might enlarge perturbations in cell length and hamper cell length stability, because cells continued to grow while cell division was inhibited [16].

We use experimental data on wild type and clock mutant cyanobacterial cells from Martins et al. 2018 [13] to determine parameter values for the gating model (Table 1). Martins et al. 2018 first entrained the circadian clock of cyanobacteria to a 12:12 LD cycle and then transferred these bacteria to a LL condition [13]. They measured doubling time, added length, birth length, and time at birth of cyanobacteria over several generations under the LL condition by single-cell tracking. We classify these wild type cells into two subpopulations based on doubling time and time at birth by applying the Gaussian mixture model following [13] (Fig. S1A–D and B). The statistics (average, standard deviation, and coefficient of variation) of birth length, doubling time, and added length for these subpopulations are listed in Table S1. We estimate the values of the added length Δ and the elongation rate *α* by using experimental data from the Δ*kaiBC* strain as Δ = 2.85 μm and *α* = log2/*τ*_0_ = 0.051 hour^−1^ (Table S1). *τ*_0_ = 13.6 hours is the average doubling time of Δ*kaiBC* strain. We set *γ*_0_ = 1 and estimate *A*, *θ_G_* and *β* by fitting the model to experimental data as described in the results section. Based on the fitting results, we use *A* = 0.91, *θ_G_* = 4*π*/3 and *β* = 50 unless otherwise mentioned.

**Table 1:** Parameter values in numerical simulation

| Parameter | Value | Unit | Description |
| --- | --- | --- | --- |
| $\tau_0$ | 13.6 | h | average doubling time of $\Delta kaiBC$ strain |
| $\alpha$ | $\ln 2/\tau_0$ | $\text{h}^{-1}$ | elongation rate |
| $\Delta$ | 2.85 | $\mu\text{m}$ | added length |
| $A$ | 0.91 | — | inhibition strength |
| $\beta$ | 50 | — | exponent of the gating function Eq. (5) |
| $\gamma_0$ | 1 | — | basal regulation |
| $\theta_G$ | $4\pi/3$ | — | gating phase when Eq. (5) takes minimum |

— : nondimension

The initial conditions of numerical simulations for Eqs. (1) and (2) are *L* = 2.85 *μ*m, *ϕ* = 0, and *θ* = 0. In the analysis of simulation data, we discard the first 100 generations to remove transient behaviors of cell length and doubling time. Then, we compute statistics for cell length and doubling time using the subsequent 101 ~ 2 × 10^4^ generations. Simulation codes for this study are written in MATLAB2020a (MathWorks). We use the ode45 solver with a fixed time step of 0.01 hours for numerical integration of the gating model Eqs. (1) and (2) (Algorithm 1).

## 3 Results

### 3.1 Bimodal distribution of doubling time by circadian gating of cell division

To examine how the circadian clock affects dynamics of cell length and doubling time, we first simulate the gating model Eqs. (1) and (2). We set *θ_G_* = 4*π*/3 in Eq. (5) so that gating occurs at subjective early night as observed in experiments. With this gating function, the circadian clock decelerates the cell phase progression near subjective early night (Fig. 3A). As a result, cell division frequency at early night is close to zero, and cells tend to divide during late night to early day when inhibition of cell division is relieved (Fig. 3A, B). Consequently, the doubling time of the cells subject to the gating increases. This increased doubling time leads to longer division lengths (Fig. 3A). In contrast, the doubling time of the cells in the subsequent generation tends to be shorter, because their long birth length increases their elongation speed Eq. (1), thereby, reducing the time required to add the length Δ. Hence, these cells divide around late day before gating occurs again (Fig. 3B), and the doubling time distribution becomes bimodal, as observed in the experimental data [13] (Fig. 3C). Thus, the effect of circadian gating appears in the doubling time of the subsequent generation through an increase in cell length.

**Figure 3:**
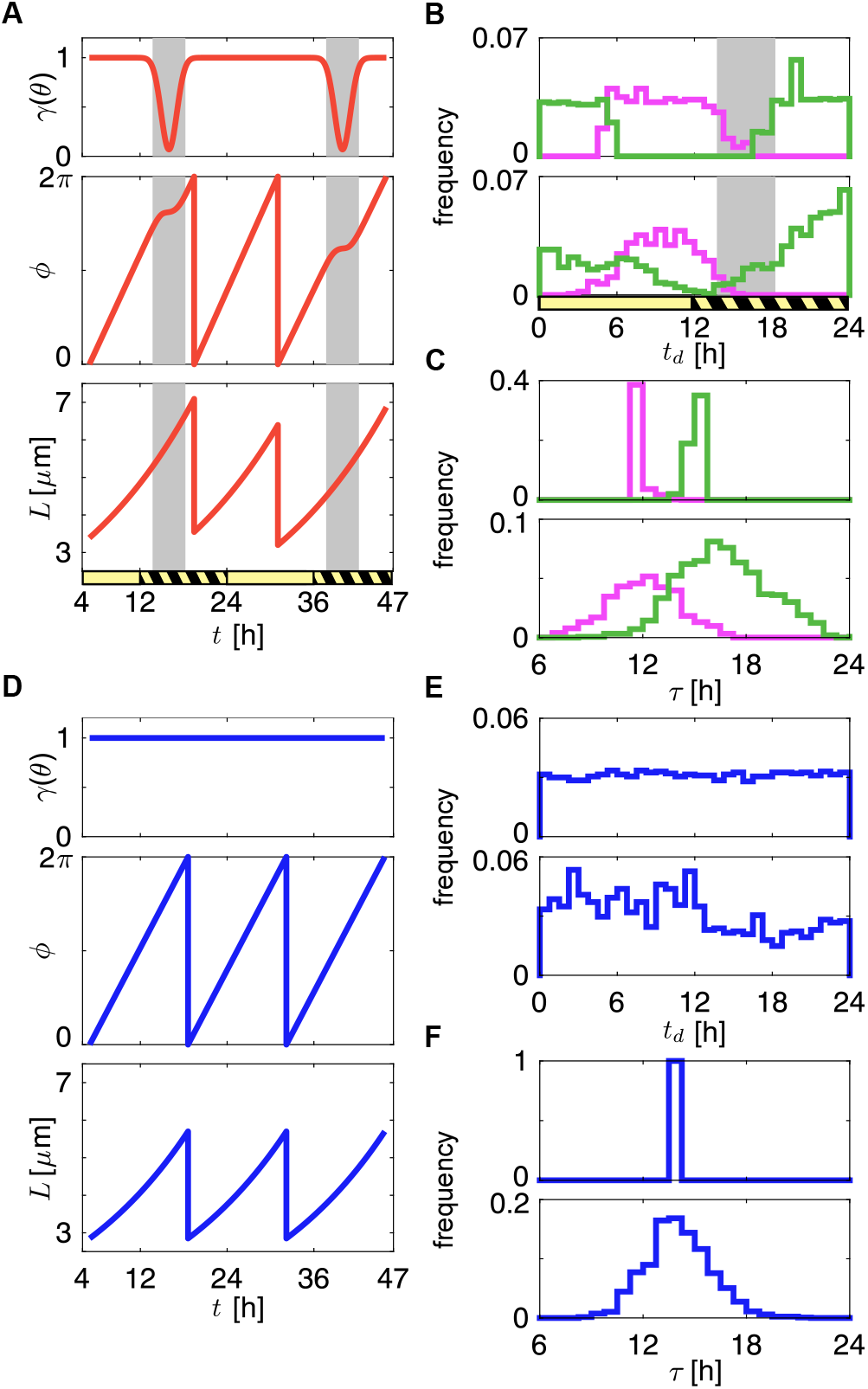
Circadian gating of cell division modulates doubling time and time at division in cyanobacteria. (A)–(C) Results of wild type. (D)–(E) Results of clock mutant. (A), (D) Time series of (top) gating function, (middle) cell phase and (bottom) cell length in simulations. (B), (E) Distributions of time at division. (C), (F) Distributions of doubling time. Top panels in (B), (C), (E) and (F) show the simulation results of the gating model and bottom panels show the experimental results of cyanobacteria under LL [13]. Green and magenta lines in (B) and (C) indicate cells whose doubling time is longer and shorter than population average (13.6 hours), respectively. The yellow and black-striped bars in (A) and (B) indicate subjective day and night, respectively. The gray shades in (A) and (B) represent circadian gating.

With the parameter set that we estimate from experimental data (see the next section), 30 cell divisions occur during 17 circadian cycles (Fig. S2A, B). Namely, cells are subject to the gating mostly once per two generations (Fig. 3A and Fig. S2C). Cells born immediately before gating give birth to a generation that is subject to gating once again (Fig. S2A). The value of cell phase *ϕ* at the gating onset changes over generations (Fig. 3A and Fig. S2A, B), indicating that cell phase is not entrained to the circadian clock.

In the absence of gating (*γ*(*θ*) = 1), each cell divides after adding the constant length Δ to its birth length through the adder mechanism. As a result, birth length *l* becomes equal to the added length, *l* = Δ (Fig. 3D). The distribution of time at division is uniform, and doubling time is *τ* = ln 2/*α* (Fig. 3E, F). These simulation results are consistent with experimental data on the circadian clock deficient Δ*kaiBC* strain [13].

To examine the effect of circadian gating on two successive generations of wild type, we introduce a delay in cell division by gating *τ_G_*. If a cell does not divide during the gating period and is fully subject to the circadian inhibition, the delay in division is obtained after integrating the gating function Eq.(5):

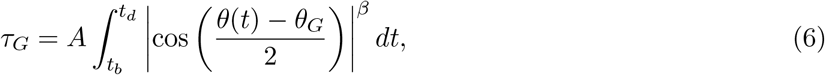

where *t_b_* and *t_d_* are the time at birth and division, respectively (see C for details). Eq. (6) indicates that the delay *τ_G_* is proportional to the inhibition strength *A*, and the proportional constant is determined by *β*. Hence, below we use this *τ_G_* to represent the inhibition strength of the circadian clock on cell division.

Figure 4 shows the dependence of doubling time, added length, and birth length on *τ_G_*. We fix *β* = 50 and determine the value of *A* in the gating model to realize each value of *τ_G_* in Fig. 4. With the increase in *τ_G_*, the doubling time of cells subject to gating increases (green triangles in Fig. 4A). In contrast, the doubling time of a cell in the generation following gating decreases as *τ_G_* increases (magenta squares in Fig. 4A), because large *τ_G_* increases its birth length and elongation rate (Fig. 3A). Therefore, the difference in doubling time between the two successive generations increases with *τ_G_*. However, the average doubling time of these two generations is independent of *τ_G_* (red circles in Fig. 4A), indicating that the doubling time in the magenta cells is shortened by the same length of time as the delay in division in the green cells.

**Figure 4:**
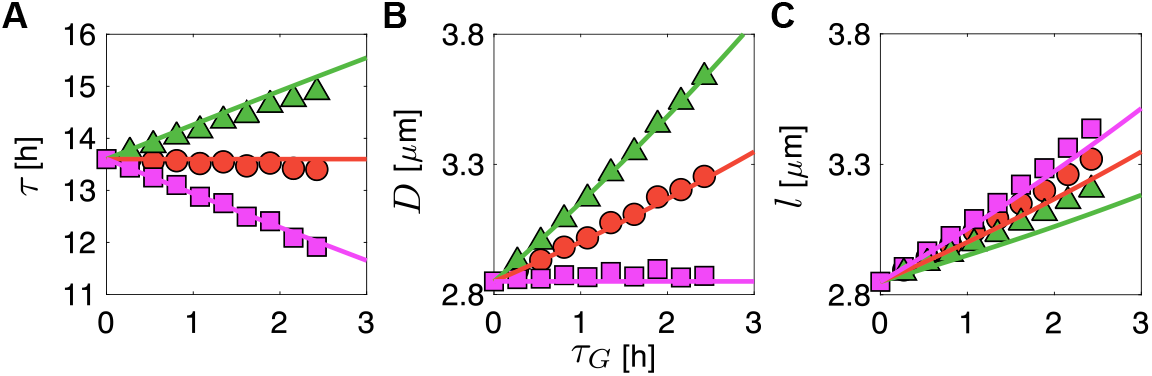
Delay in cell division *τ_G_* caused by circadian gating influences doubling time and cell length. Dependence of (A) doubling time, (B) added length and (C) birth length on *τ_G_*. In (A)–(C), green indicates results for cells subject to circadian gating. Magenta in (A)–(C) indicates results for cells not subject to gating. Red indicates the average of these two generations. Marks and lines indicate the results of the gating model and the two-state model, respectively.

The added length of cells subject to the circadian gating is longer than Δ because of the delay in division (green triangles in Fig. 4B). Accordingly, the added length of this generation increases with the delay *τ_G_*. In contrast, the added length of cells in the subsequent generation that divides before the gating is independent of *τ_G_* and is nearly Δ (magenta squares in Fig. 4B). As a result, the average added length over the two successive generations increases with *τ_G_* (red circles in Fig. 4B).

The long added length with large *τ_G_* in cells subject to gating results in long birth length of the subsequent generation (magenta squares in Fig. 4C). Without being affected by the circadian clock, these magenta cells divide after adding Δ to their long birth length. Hence, the birth length of cells subject to gating also increases with *τ_G_* (green triangles in Fig. 4C). We notice that the dependence of the average added and birth lengths on *τ_G_* are almost the same, as indicated by red circles in Fig. 4B and C. We will return to this point later in the results section. In summary, our analysis indicates that the circadian gating of cell division influences both doubling time and cell length in each generation.

### 3.2 Estimation of delay in cell division by circadian gating

We then estimate the delay *τ_G_* in cell division by circadian gating in wild type cyanobacteria under LL by fitting the gating model to experimental data from Martins et al. 2018 [13]. Specifically, we compare the doubling time *τ*, added length *D* and birth length *l* of wild type cells born at *t_b_* from our simulation data with experimental data [13] using the least-squares method (see D for details; Fig. 5). We first determine the gating phase *θ_G_* = 4*π*/3 based on the square error between simulation and experimental data (Fig. S3A–F). Figure 5A shows the dependence of the square error on *A* and *β* with *θ_G_* = 4*π*/3 in the gating function Eq. (5). There is a valley of square error minima in the *A-β* space, as indicated by gray triangles in Fig. 5A. We find that the delay *τ_G_* is almost constant, *τ_G_* = 2.44 ± 0.0714 hours along this valley (Fig. 5B). To retain *τ_G_* within the above range, a small *A* (i.e., weak inhibition) requires a small *β* (i.e., long gating duration). However, quite a few cells divide during gating with weak inhibition of cell division (small *A*). Therefore, we set *A* = 0.91 in the subsequent analyses to prevent cell divisions during gating as observed in experimental data (Fig. 3B).

**Figure 5:**
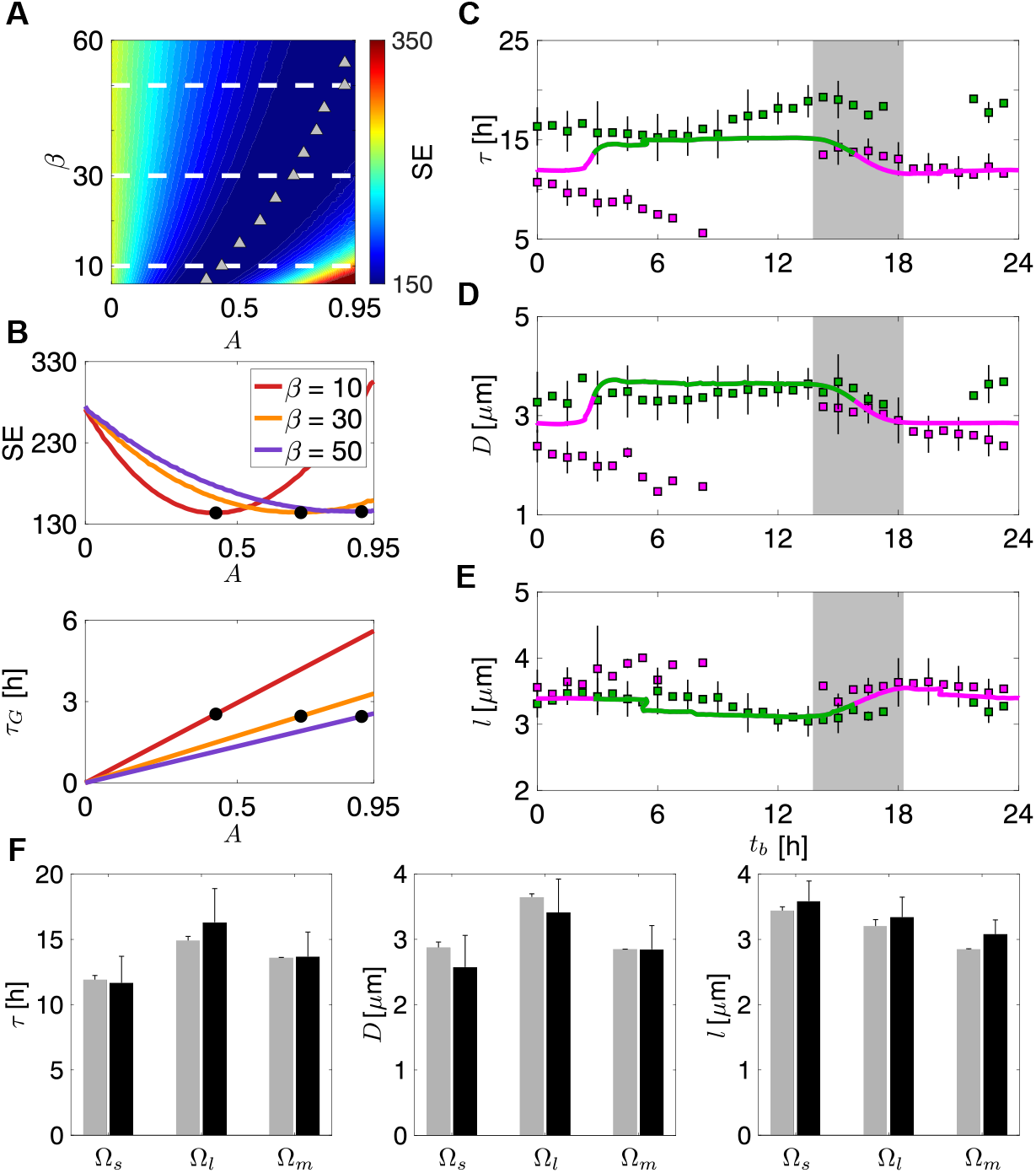
Circadian gating causes 2.45-hour delay in cell division. (A) Dependence of square error (SE) on *A* and *β* in the least-squares method. The color indicates the value of SE between simulations and experimental data from [13]. White broken lines indicate *β* slices of which SE values are plotted in (B). Gray triangles indicate the values of *A* where SE takes a minimum along each *β* slice. (B) (top) SE as a function of *A* for *β* = 10, 30, 50. (bottom) *τ_G_* as a function of *A* for *β* = 10, 30, 50. Black dots indicate minimum values of SE for each *A* and *β*. (C)–(E) Dependence of (C) doubling time *τ*, (D) added length *D*, and (E) birth length *l* on time at birth *t_b_* in experiment (squares) and simulation (lines) with the best fitted parameter set: *A* = 0.91, *β* = 50 and *θ_G_* = 4*π*/3. Green and magenta indicate cells with longer and shorter doubling time than population average, respectively. Squares and error bars indicate average and standard deviation for experimental data, respectively. The average and standard deviation are computed with a bin size of 0.75 hours. Gray shades in (C)–(E) indicate circadian gating. (F) Average doubling time (left), added length (middle), and birth length (right) for each cell type in experiment (gray) and simulation (black). Ω_*s*_ and Ω_*l*_ represent cells with a shorter and longer doubling time than population average, respectively. Ω_*m*_ represents Δ*kaiBC* strain.

The fitted model with *τ_G_* = 2.45 captures the overall trends in the dependence of *τ*, *D*, and *l* on *t_b_* (Fig. 5C–E). The fitted lines for *τ* and *l* include small kinks or folds at around *t_b_* ≈ 5.4 and 20 where the same *t_b_* provides three different values for *τ* or *l* (Fig. 5C, E). The cells in the kink at t_b_ ≈ 5.4 are generated by the divisions of their mother cells for which *t_b_* is between 15 and 18 (gray shades in Fig. 5C–E). The doubling time of the mother cells decreases with *t_b_* within the gating window in the form of a S-shaped curve (Fig. 5C). Because the time at division of these mother cells is expressed as *t_d_* = *t_b_* + *τ*, decreases in *τ* with increases in *t_b_* allows a cell born earlier and another cell born later to divide at the same time. Since the added lengths of such cells are different (Fig. 5D, E), the birth length and doubling time of their daughter cells are also different, causing the folding structure at *t_b_* ≈ 5.4. Division of cells that emerge at *t_b_* ≈ 5.4 results in the other fold at *t_b_* ≈ 20.

The gating model does not reproduce the long doubling time at *t_b_* = 10 ~ 16 (green squares in Fig. 5C) and the long birth lengths at *t_b_* = 0 ~ 8 (magenta squares in Fig. 5E) observed in experimental data. These long doubling time and birth lengths may be interrelated, because the long doubling time of the green cells in Fig. 5C can result in their long division lengths and, therefore, a long birth length of their daughter cells (Fig. 5E). These deviations of the current gating model from experimental data suggest that although the current model assumes a constant value for inhibition strength *A*, its value varies with cell phase *ϕ* in living cyanobacteria. In addition, the gating model does not reproduce the short added length of cells at *t_b_* = 0 ~ 8 (magenta squares in Fig. 5D). These deviations in added length may be partly due to the simplifying assumption of the constant added length Δ in the current gating model.

We also confirm that the fitted model reproduces the average doubling time, added length, and birth length of each cell type that is observed in experiments (Fig. 5F and Table S1). In summary, the gating model captures similar statistical trends as is observed in the experimental data. The circadian clock delays cell division about 2.45 hours at early subjective night.

### 3.3 Simplification of the gating model by averaging cell phase dynamics

Although the best fitted values of the inhibition strength *A* and exponent *β* in the gating function Eq. (5) were not determined uniquely, we estimated the delay *τ_G_* in the previous section. To analyze cell length stability based on the estimated value *τ_G_* = 2.45, here we introduce a simplified description of the circadian gating that does not include un-estimated parameters *A* and *β*, but includes estimated *τ_G_*. Along with the calculation of cell length stability, we also derive equations for the average birth length, added length, and doubling time (Fig. 4).

In the gating model Eqs. (1) and (2), only a few cells divide within the time window where cell phase is slowed by the circadian clock. In the simplified description, we neglect such cells and only consider those that do not divide during the circadian gating. In addition, we assume that the subsequent generation is unaffected by the gating. This simplification is based on the clear bimodality of the simulated doubling time distribution (Fig. 3C).

Let *n* specify the generation of cells (*n* = 0, 1, 2,…). We assume that cells in even generations are subject to the deceleration of cell phase by the circadian clock, and refer to these cells as the inhibited generation. Cells in odd generations are not influenced by the clock and divide when their added length becomes Δ. We refer to these cells as the uninhibited generation. Then, the time derivative of cell phase is described as:

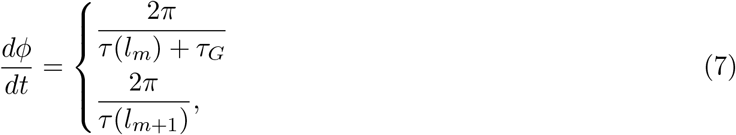

where *m* = 0, 2, 4,… and *τ*(*l_x_*) = ln(1 + Δ/*l_x_*)/*α* is the doubling time when the generation *x* is not influenced by the circadian clock. *τ_G_* is the delay in cell division due to the circadian gating described in Eq. (6). We use the estimated value *τ_G_* = 2.45 hours for the following analyses. In this two-state model Eq. (7), we average the cell phase progression of the inhibited generation over time, and approximate cell phase in the gating model by the obtained average speed between its birth and subsequent division (Fig. 6A and see C for details). Consequently, the cell phase of the inhibited generation increases at a constant, but slower, rate compared to that of the uninhibited generation (Fig. 6B). Although this constant deceleration changes the trajectory of the cell phase from those in the gating model, doubling time remains the same between these two models (Fig. 6A). Eq. (7) implies that the division of those cells subject to circadian gating is determined by the combination of the adder mechanism and a timer mechanism [1] where the cell has to wait a constant time *τ_G_* after *τ*(*l*) to divide.

**Figure 6:**
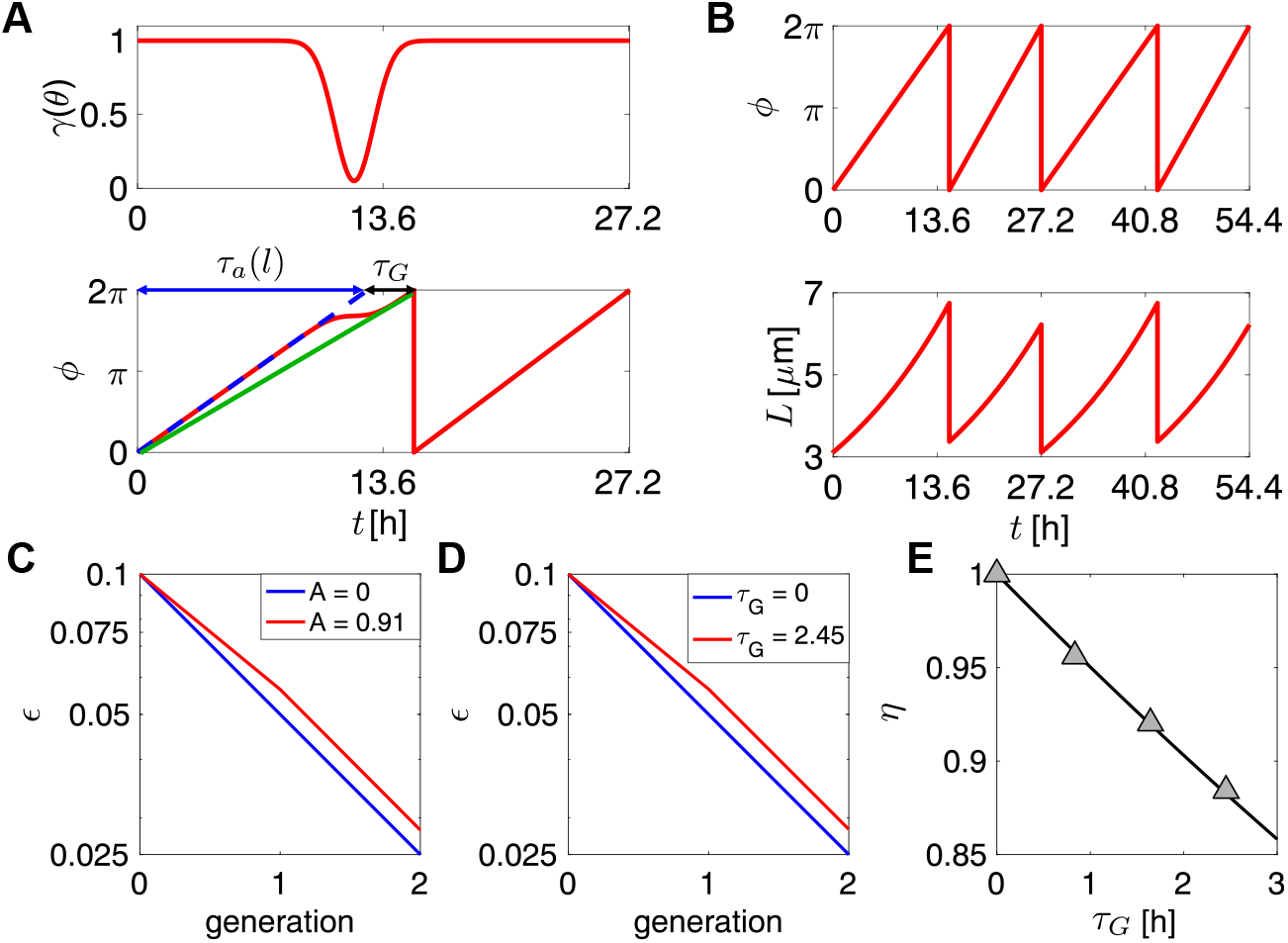
The decay rate of cell length perturbation decreases with the delay in cell division *τ_G_* in both gating and two-state models. (A) Averaging of cell phase speed. Time series of (top) the gating function Eq. (5) and (bottom) cell phase. Blue broken line indicates cell phase in the absence of circadian gating. The red line indicates cell phase modulated by the gating function shown in the top panel. The green line indicates cell phase at the averaged speed. (B) Time series of (top) cell phase and (bottom) cell length in the two-state model. (C), (D) Time series of perturbation in cell length *∊* added in the zeroth generation in (C) gating model and (D) two-state model. Blue lines in (C) and (D) indicate results for *A* = 0 and *τ_G_* = 0, respectively. Red lines in (C) and (D) indicate results for *A* = −0.91 and *τ_G_* = 2.45, respectively. The vertical axis is in log scale. (E) Dependence of relative decay rate *η* on *τ_G_*. Gray triangles indicate results of the gating model. Line indicates the result of the two-state model η = 2^−*τ_G_*/*τ*_0_^.

We describe the time evolution of cell length by Eq. (1) and simulate this two-state model Eqs. (1) and (7) to confirm that it captures the key properties of the gating model (Fig. 6B). We use the ode45 solver in MATLAB for numerical integration. Starting from an initial birth length for generation 0, the system converges to a periodic solution (Fig. 6B). Due to the model’s assumptions, the doubling time of two successive generations differs. The added length of the inhibited generation is longer than that of the uninhibited generation. As a consequence, the birth length of the uninhibited generation is longer than that of the inhibited generation. Thus, the periodic solution in the two-state model Eqs. (1) and (7) captures the qualitative properties of the gating model.

### 3.4 Dependence of cell length and doubling time on delay *τ_G_* in the two-state model

Using the 2-state model Eqs. (1) and (7), we derive the equations for birth length, added length, and doubling time. We start with difference equations for the birth length for the uninhibited and subsequent inhibited generations:

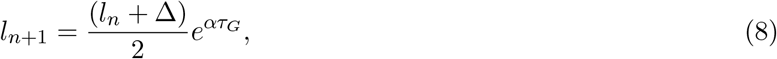

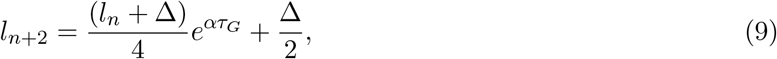

where *l_n_* is the birth length of the inhibited generation *n* (*n* = 0, 2, 4,…). After the convergence to the period-2 solution, the birth lengths of the uninhibited and inhibited generations, *l_s_* and *l_l_*, respectively, can be written as:

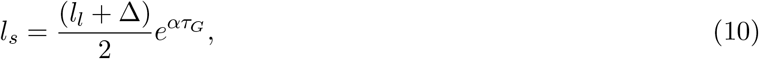

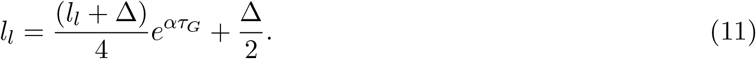

By solving Eqs. (10) and (11), we obtain:

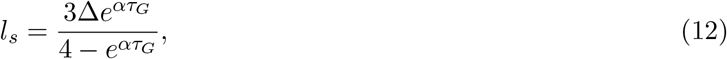

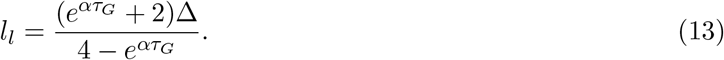

Eqs. (12) and (13) indicate that the periodic solution exists only when *ατ_G_* < ln 4. Both Eqs. (12) and (13) include *τ_G_*, indicating the inheritance of gating effect by the subsequent generation. *l_s_* and *l_l_* increase with the increase in *τ_G_*, which is consistent with the result of the gating model (Fig. 4C). Furthermore, using Eqs. (12) and (13), the average birth length 〈*l*〉 can be written as:

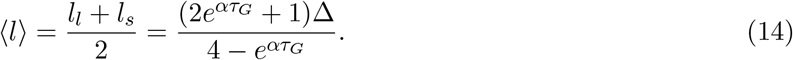

The added length Δ_*l*_ for the inhibited generation is the difference between the division length *L_l_* = 2*l_s_* and birth length *l_l_*:

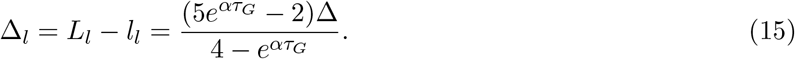

Eq. (15) indicates that the added length of the inhibited generation increases with *τ_G_* as in the gating model (Fig. 4B). The added length of the uninhibited generation is Δ_*s*_ = Δ. Hence, their average 〈Δ〉 is:

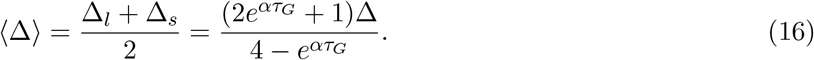

Thus, 〈*l*〉 in Eq. (14) and 〈Δ〉 in Eq. (16) are equivalent and explain the simulation results in the gating model (Fig. 4B, C). Furthermore, from *l_l_* and *L*l** = 2*l_s_*, we define change in added length *δ* due to inhibition as:

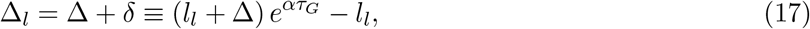

where

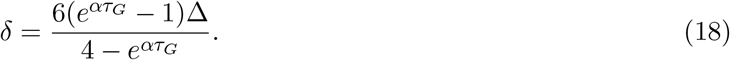

By using Eqs. (12), (13) and (17), we obtain the ratio between the differences in added lengths and birth lengths (Δ_*s*_ – Δ_*l*_)/(*l_s_* – *l_l_*) as – *δ*/(*l_s_* – *l_l_*) = −3.0 by substituting the estimated values of *α*, Δ and *τ_G_* (Table 1). Consistently, the corresponding ratio calculated from the experimental data for average added lengths and birth lengths (Table S1) is −3.40. Thus, circadian gating causes a negative correlation between added length and birth length. This contrasts with the independence of added length from birth length in the original adder mechanism. Using the above defined *δ*, doubling time of the inhibited generation *τ_l_* can be expressed as:

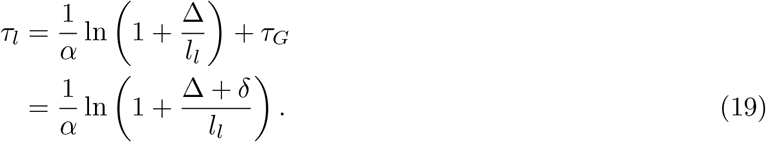

Eq. (19) indicates that the doubling time of cells subject to circadian gating is determined by the adder mechanism with a modulated added length Δ + *δ* in the period-2 solution. In addition, because the added length of the uninhibited generation is Δ, its doubling time is τ_*s*_ = *α*^−1^ ln(1 + Δ/*l_s_*). Thus, after the convergence to the period-2 solution, the system alternates the adder mechanism with different added lengths Δ_*l*_ and Δ.

Finally, using Eqs. (12), (13) and (19), we obtain the doubling time of the inhibited and uninhibited generations in the period-2 solution:

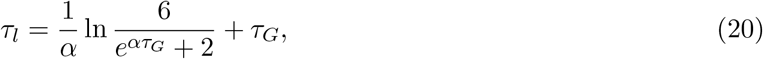

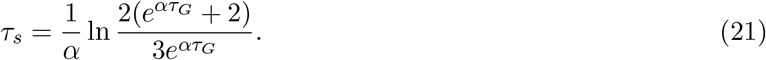

Eq. (21) indicates that the change in *τ_G_* influences doubling time of uninhibited generation. The doubling time is approximately proportional to *α*^−1^ and more sensitive to the elongation rate *α* than *τ_G_*. Finally, the average doubling time is:

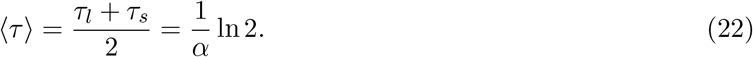

Thus, the average doubling time of the uninhibited and inhibited generations is independent of *τ_G_* and only depends on the elongation rate *α*. We have also shown the independence of *τ_G_* in the gating model (Fig. 4A). From Eq. (22), *τ_s_* can be expressed as:

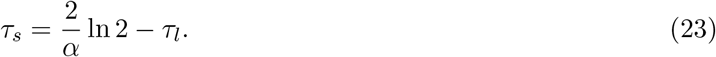

The circadian gating lengthens doubling time of the inhibited generations, but equally shortens that of the uninhibited generations by increasing birth length. In summary, the two-state model explains the numerical results of the gating model.

### 3.5 Cell length stability

Eq. (15) indicates that the added length due to the circadian gating Δ_l_ becomes longer than that due to the adder mechanism alone (i.e., Δ_*l*_ > Δ). This implies that circadian gating might also enlarge perturbations in cell length caused by physiological fluctuations. Here we compare the decay rate of cell length perturbations in the presence of circadian gating with that of the adder mechanism alone.

To derive the decay rate in the two-state model, we perform stability analysis of the period-2 solution. We add a perturbation *∊_n_* to the birth length of an inhibited generation *n* as *l_n_* = *l_l_* + *∊_n_*. Then, the perturbation in the subsequent uninhibited generation can be written from Eq. (9) as:

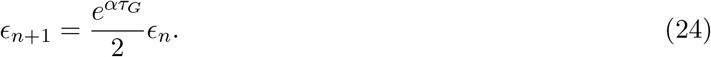

The factor *e^ατ_G_^* represents the enlargement of the perturbation due to the circadian gating. Using the relation *ατ_G_* = *τ_G_* ln 2/*τ*_0_, we write Eq. (24) as:

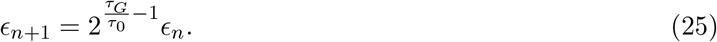

Eq. (25) shows that when *τ_G_* is much smaller than *τ*_0_ (*τ_G_*/*τ*_0_ ≪ 1), the effect of the circadian clock on cell length stability becomes negligible, as (*τ_G_*/*τ*_0_) – 1 ≈ −1. In this limit, cell length perturbation decays as it does in bacterial cells with the adder mechanism alone. With values of *τ_G_* = 2.45 and *τ*_0_ = 13.6 estimated from experimental data, we find *τ_G_*/*τ*_0_ = 0.18 for the cyanobacteria. Hence, Eq. (25) predicts that the decay rate of cell length perturbation in cyanobacteria is 2^0.82^/2 = 0.88 times smaller than other bacteria with the adder mechanism alone.

Because the generation *n* + 1 is not subject to circadian regulation, we can write *∊*_*n*+2_ based on the adder mechanism:

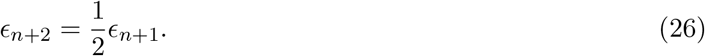

By substituting Eq. (24) into Eq. (26), we obtain:

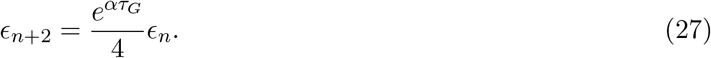

Thus, as long as the period-2 solution exists *e^ατ_G_^* < 4, the solution is stable, ensuring cell size homeostasis.

Finally, we confirm the above calculations by simulating the original gating model Eqs. (1) and (2). We increase the birth length of the inhibited generation by 10% from its average value in the gating model, and perform numerical simulations to examine the decay of the perturbation *∊_n_*. We define the decay rate of perturbation relative to that of the adder mechanism as *η* = (*∊_n_*/*∊*_*n*+1_)/2 for the gating model. Without circadian gating (*A* = 0), the perturbation in cell length halves after division due to the adder mechanism (blue line in Fig. 6C). In the presence of circadian gating with an estimated value of *A* (*A* = 0.91), perturbation decays slower (red line in Fig. 6C). We observe similar behavior in the two-state model where *τ_G_* = 2.45 that corresponds to *A* = 0.91 in the gating model (Fig. 6D). In simulations of both gating and two-state models, the decay rates in the presence of circadian gating is 0.87 times smaller than its absence (Fig. 6C–E), confirming the analytical calculation from the two-state model Eq. (25). The relative decay rate η further decreases as the delay in cell division *τ_G_* increases (Fig. 6E). Thus, stronger gating indeed reduces the speed at which cell length returns to its original value after being perturbed.

In summary, circadian gating decreases the decay rate of cell length perturbation by extending the doubling time. However, with the estimated gating parameters in cyanobacteria, the influence of the gating on the decay rate is negligible. This is because doubling time determined only by cell length is sufficiently longer than the delay in cell division by gating. Thus, the adder mechanism in cyanobacteria ensures cell size homeostasis under LL.

## 4 Discussion

Circadian control of cell division was first observed in cyanobacteria under LL [12], and later found in eukaryotes including mammalian cells [22–26]. Mori et al. 1996 proposed that the circadian clock gates cell division at early subjective night [12]. This hypothesis was mathematically formulated by Yang et al. 2010 [15]. Later studies examined cyanobacterial cell division using mathematical modeling together with single-cell imaging [10, 13, 16, 20]. However, it has remained unclear how cyanobacteria maintain their characteristic cell length over many generations in the presence of circadian regulation of cell division. To address this question, we combined previous modeling approaches [15, 21] to describe cell division regulated by both the adder mechanism and the circadian clock. We clarified the effects of circadian gating on cell length, doubling time, and cell length stability in cyanobacteria.

Under LD cycles, cyanobacterial cells elongate and divide only in the presence of light [12, 13]. In previous experiments [12, 13], cyanobacteria were first entrained to a 12-hour light and 12-hour dark cycle, then transferred to LL conditions to measure the cell division frequency. Since cells continue to divide in the presence of light, circadian gating of cell division is apparent under LL [12, 13, 15]. We and other researchers have consistently identified that circadian gating occurs at the early subjective night [12, 13, 15]. However, the timing of circadian gating might depend on the parameters of LD cycles that cyanobacteria were exposed to. For example, the day length of a LD cycle, such as 16-hour light and 8-hour dark, changes the time at which KaiC is maximally phosphorylated [27]. In addition, the period of a LD cycle may also change the phase of entrainment of KaiC phosphorylation rhythms. Thus, through affecting cyanobacterial circadian clock, LD cycles could shift the timing of gating. Characterizing the effect of LD cycles on gating would be crucial for understanding cell division across a variety of day lengths, e.g., in different seasons, as well as the evolution of cyanobacteria as the rotation speed of ancient earth decreased over time (thus increasing day length) [28, 29].

Our model predicts that a cell with a longer doubling time due to gating gives birth to elongated daughter cells, and that these daughter cells elongate faster and divide during subjective day. Furthermore, our calculation suggests that the average doubling time of a cell lineage is the same as that of the Δ*kaiBC* strain. We found a nearly one hour difference between the average doubling time of wild type (14.7 hours) and Δ*kaiBC* (13.6 hours) strains in the experimental data [13]. More detailed validation of these predictions requires cell lineage data for many generations by cell-tracking algorithms [30–33]. Such lineage analyses will also reveal the relationship between the doubling time of single cyanobacteria and that of whole populations, as has been done for *E. coli* [33]. In *E. coli*, the population doubling time is always shorter than the average doubling time of single cells [33]. Hence, it would be an interesting future work to study how the variation of doubling time caused by circadian gating in single cyanobacteria influences the population doubling time.

Cell size homeostasis in bacterial species that do not possess a circadian clock, such as *E. coli* and *B. subtilis*, has been studied by both experimental and theoretical approaches [1, 6]. These bacterial cells divide after adding a constant size to their birth size [1]. In this adder mechanism, perturbation in cell length decays at a rate of 2^−1^ every division [6]. In this study, we revealed that the decay rate of perturbation in cells subject to gating is determined by the ratio of delay in cell division by gating *τ_G_* to the average doubling time *τ*_0_ as 2^*τ_G_*/*τ*_0_-1^. We estimated *τ*_0_ = 13.6 hours from the average doubling time of Δ*kaiBC* strain [13], and *τ_G_* = 2.45 hours by fitting the gating model with a least-squares method. The estimated ratio *τ_G_*/*τ*_0_ is smaller than 0.2, indicating that perturbation decays at almost the same speed as it does in the adder mechanism. In other words, because the average doubling time is proportional to the inverse of the elongation rate *τ*_0_ ∝ 1/*α*, our results indicate that the slow elongation of cyanobacteria prevents extension of cell length perturbation when the circadian clock inhibits cell division. Our prediction can be validated experimentally by monitoring changes in cell length after transient cell elongation. Currently, mutant strains or chemical treatments that alter *τ_G_* are not available. In contrast, a previous imaging study showed that *τ*_0_ depends on light intensity [15]. The same study also suggested that light intensity does not significantly alter circadian gating by using mathematical modeling [15]. Thus, light intensity may significantly change only *τ*_0_ and modulate the ratio *τ_G_*/*τ*_0_. Perturbation in cell length may be realized by the transient elevation of KaiC ATPase activity, which inhibits proper localization of FtsZ at midcell [16]. For example, overexpression of KaiA from IPTG-inducible *kaiA* gene by IPTG addition resulted in elevated KaiC ATPase activity that inhibited cell division and caused cell elongation [16]. Then, KaiA protein levels were reduced back to physiological levels by IPTG removal. Thus, transient perturbation of cell length by overexpression of KaiA would validate current theoretical predictions for cell length stability in cyanobacteria.

Based on statistical modeling with single-cell tracking data, Martins et al. 2018 [13] recently proposed that the cyanobacterial circadian clock transiently promotes cell division at late subjective day. During the rest of day, the circadian clock reduces the probability of cell division [13]. Thus, the model by Martins et al. 2018 includes two opposite influences of the circadian clock on cell division. Cyanobacterial genes that promote cell division have not been identified yet. In *E. coli*, however, overexpression of several genes results in the promotion of cell division. For example, *nrdAB* can enhance dNTP synthesis, and its overexpression shortens the time between the initiation and the completion of DNA replication [34]. In contrast, *ftsZA* promotes assembly of the divisome that constricts cell membranes during division [35], and overexpression of *ftsZA* decreases the time between the completion of DNA replication and that of cell division. Indeed, the overexpression of these two genes by optgenetics led to the promotion of cell division [34]. If the cyanobacterial circadian clock induces specific expression of these genes, it would promote cell division, as Martins et al. 2018 proposed.

In contrast, the gating model we have adopted assumes that the circadian clock transiently inhibits division only at early subjective night. We showed that this simple gating model can reproduce most of the statistics for doubling time and cell length. This is because there are a number of similarities between the model by Martins et al. 2018 and ours. First, both models include the adder or adder-like mechanisms to describe cell length control. Second, both models include time windows where the circadian clock inhibits cell division. The inhibition of cell division by the circadian clock together with the adder mechanism increases cell length at division. Hence, it explains longer birth length and added length in the wild type than in the Δ*kaiBC* strain under LL in experiments [13]. Third and last, both models include a transient change in the effect of circadian clock on cell division. Such a transient change results in temporal variation of the cell division frequency and different doubling times of cells, as observed in the wild type. These three common aspects, therefore, would be essential for the description of cell division in cyanobacteria.

In this study, we adopted a phenomenological model of the circadian gating. Although the current model only considers the influence of the circadian clock on cell division, cell division could also affect the phase of the circadian clock [22, 36, 37]. For example, because gene copy number increases during DNA replication, cell divisions can cause temporal changes in the transcription rate of *kaiABC* genes and shift the phase of the circadian clock [37]. A possible future work, therefore, would be to develop a mathematical model that includes molecular details of KaiABC oscillators, the adder mechanism, and clock-dependent cell division control. The mechanism for the autonomous phosphorylation cycle of KaiC protein has been studied extensively by both experimental and theoretical approaches [38]. The phosphorylation state of KaiC is associated with its ATPase activity [19]. The precise mechanisms behind circadian gating are not yet known, but Dong et al. 2010 proposed that elevated KaiC ATPase activity activates RpaA through SasA, and then activated RpaA inhibits the localization of FtsZ in the midcell, ultimately leading to the inhibition of cell division [16]. Inspired by the necessity of FtsZ accumulation for cell division, Ho et al. 2020 proposed a model that cyanobacterial cell division occurs when the concentration of a protein, termed divisor, reaches a certain threshold [20]. If the production rate of this divisor protein is proportional to the elongation speed of individual cells, these cells divide following the adder mechanism. Ho et al. 2020 further assumed that the circadian clock modulates the production rate of the divisor protein. It is still unclear whether the FtsZ protein works as a divisor protein, because its overexpression inhibited cell division and produced filamentous cells that were much longer than the wild type [39]. It is thus likely that both appropriate amount and localization of FtsZ at the midcell are important factors for cell division [16]. Besides, other growth and division mutants with various morphological phenotypes have been discovered [40]. Some of these mutated genes may be regulated by the circadian clock. In addition, the regulation of cyanobacterial DNA replication by the circadian clock [41] may be one of the composing elements of gating. Thus, although some key aspects remain to be revealed, the construction of a mathematical model based on molecular mechanisms for cyanobacterial cell division would be within reach in the near future.

Circadian control of cell division has been observed not only in cyanobacteria but also in fish and mammals [22, 23, 25, 26, 42]. In mammals, the transcription of genes involved in the cell cycle, such as *c-Myc* and G2/M inhibitor *Wee1*, is regulated by circadian clock proteins [24, 25, 43]. Mammalian circadian clock protein PER2 interacts with the tumor suppressor P53 protein and prevents its degradation [26]. Furthermore, the proliferation rate of B16 melanoma cells is reduced by the circadian clock. In addition, it has been shown that the division of eukaryotic cells, including budding yeast [4] and several mammalian cells [44], is regulated by the adder mechanism. Hence, the framework developed in the current study would also be applicable to the analysis of cell size homeostasis in these eukaryotic cells.

## Acknowledgment

We would like to thank the members of Chronogenomics Laboratory at Kanazawa University for useful discussion. This work was supported by the JSPS KAKENHI grant number 19H04955, 19H04772 and 20K06653 to KU.

## Author contributions

YK: Conceptualization, Methodology, Software, Formal analysis, Investigation, Visualization, Writing - Original Draft, Writing - Review & Editing.

HT: Conceptualization, Writing - Review & Editing, Supervision.

KU: Conceptualization, Methodology, Formal analysis, Writing - Original Draft, Writing - Review & Editing, Supervision, Funding acquisition.

## Appendix A Adder mechanism

We review the doubling time and cell length stability of the adder mechanism [6, 21]. From Eq. (1), the cell length *L* at time *t* is *L*(*t*) = *le^αt^* where *l* is the cell length at birth (birth length). In the adder mechanism, cells divide after adding a constant length Δ to its birth length. Hence, doubling time *τ* satisfies Δ = *le^ατ^* – *l*. By solving this equation with respect to *τ*, we obtain

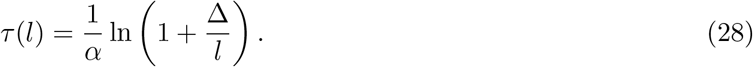

Thus, the doubling time depends on the birth length, elongation rate, and added length in the adder mechanism.

To see the stability of cell length, we consider the difference equation of the birth length for the generation *n* + 1:

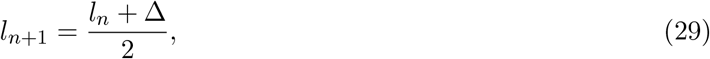

where we assume that cell length becomes exactly half after division. The steady state of the difference equation *l*\* is *l*\* = Δ. We then consider a perturbation from the steady state *l_n_* = *l*\* + *∊_n_*. By substituting this equation into Eq. (29), we obtain

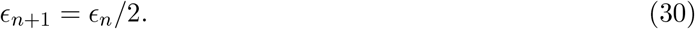

Thus, perturbation in cell length decays at the rate of 2^−1^ per division in the adder mechanism.

## Appendix B Classification of cells with the Gaussian mixture model

We use the Gaussian mixture model (GMM) in MATLAB (fitgmdist) to classify cyanobacterial cells from the experimental data by Martins et al. 2018 [13] into two groups based on doubling time and time at birth. To deal with the 24-hour periodicity of time at birth, we triplicate the original 24-hour data (Fig. S1A). The time at birth in the triplicated data ranges from −24 to 48 hours. Then, the GMM algorithm is initialized with *k*-means clustering with the cluster number of 6 (*k* = 6). After the application of the GMM, we further classify cells into two groups (long and short doubling time) by comparing their likelihoods. Specifically, if a cell attains the largest likelihood within clusters 2 or 4 (Fig. S1A), we assign the cell to the long doubling time group. In contrast, if a cell attains the largest likelihood within the cluster 3, it is assigned to the short doubling time group. These procedures classify cells into two subpopulations based on doubling time and time at birth, consistent with the classification by Martins et al. 2018 (Fig. S1B, C) [13]. In the original data, circadian time (CT) from 0 to 12 represents subjective night, and CT from 12 to 24 represents subjective day (Fig. S1B) [13]. To follow the convention in the circadian field (i.e., CT 0–12 as subjective day and CT 12–24 as night), we shift time at birth and division by 12 hours (Fig. S1D).

## Appendix C Delay in cell division by circadian gating

We derive the delay in cell division caused by the circadian gating *τ_G_*. Without loss of generality, we assume *θ_G_* = 0 in the gating function Eq. (5). We consider a cell that is born at time *t_b_*, is fully subject to gating, and divides at time *t_d_*. Integration of Eq. (2) with respect to *t* leads to

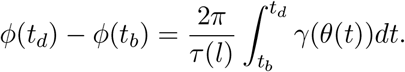

Because the cell divides at time *t_d_*, we write the above equation as

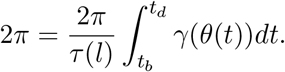

By substituting Eq. (5) with *γ*_0_ = 1, we obtain

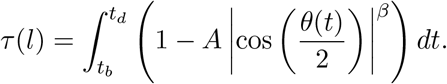

Because *t_d_* – *t_b_* = *τ*(*l*) + *τ_G_*, the delay *τ_G_* can be written as

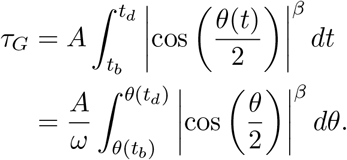

where we used Eq. (4). Note that | cos(*θ*/2)|^*β*^ = 0 for *θ* = (2*n* + 1) *π*, and | cos(*θ*/2)|^*β*^ = 1 for *θ* = 2n*π* (*n* = 0, 1, 2,…). Hence, if a cell is subject to gating and its doubling time is shorter than the circadian period, (2*n* – 1)*π* < *θ*(*t_b_*) < *θ*(*t_d_*) < (2*n* + 1)*π*. Furthermore, if *β* is sufficiently large, | cos(*θ*/2)|^*β*^ ≈ 0 except for the vicinity of *θ* = 2*nπ* (Fig. 2E). Hence, we replace the range of integration as

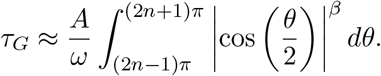

If *β* is an integer number, we obtain

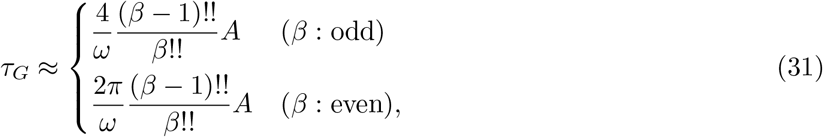

where !! represents double factorial.

## Appendix D Estimation of *τ_G_* by a least-squares method

To estimate *τ_G_*, we fit the gating model to the experimental data by minimizing the square error between experimental and simulation data. The dataset from Martins et al. 2018 [13] includes information about doubling time *τ*, added length *D*, and birth length *l* of cells with time at birth *t_b_*. The sampling interval of their data is Δ*t_s_* = 0.75 hours. For each time point *t_j_* = *j*Δ*t_s_* (*j* = 0, 1,…, *T* and *t_T_* = 24), there are experimental data for *N_t_j__* cells. We define square error at time point *t_j_* as

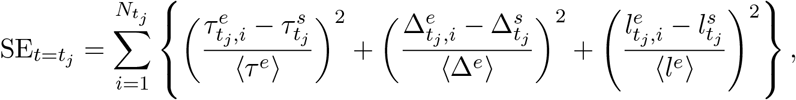

where 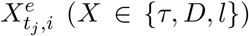 (*X* ∈ {*τ*, *D*, *l*}) represents the experimental data for cell *i* with time at birth *t_j_*, and 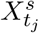 represents the simulation data for the cell with time at birth *t_j_*. Each square error is normalized by the average doubling time 〈*τ^e^*〉, added length 〈Δ^*e*^〉, and birth length 〈*l^e^*〉 of experimental data. In simulations, we may not be able to sample the cell that is born exactly at *t_j_*. In such cases, we interpolate 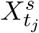 linearly using sampled simulation data at two nearest neighboring time points of *t_j_*, 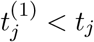 and 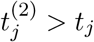. Then, we calculate the total square error over all time points:

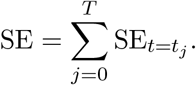

Fig. S3A–F shows the dependence of SE on *A* and *β* in the gating function Eq. (5) for different values of the gating phase *θ_G_*. We find SE minima in the *A-β* plane with *θ_G_* = 4*π*/3 (Fig. S3E). Therefore, we further search *A* and *β* values that minimize SE by fixing *θ_G_* = 4*π*/3 (Fig. 5A).

