## Supplemental Figure for "Cell size homeostasis under the circadian regulation of cell division in cyanobacteria"

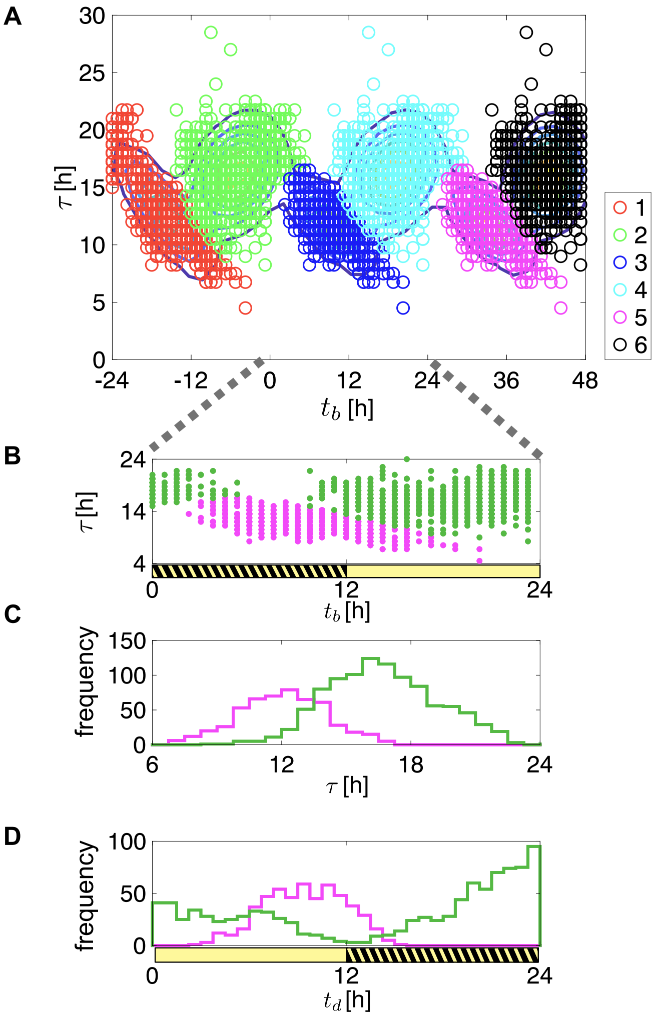


Figure S1: Classification of cells by the Gaussian mixture model (GMM) based on time at birth $t_b$ and doubling time $\tau$. (A) Dependence of $\tau$ on $t_b$. The experimental data between $t_b = 0$ and $t_b = 24$ hours are triplicated to deal with the 24-hour periodicity of $t_b$ in GMM. Colored circles indicate the six clusters obtained by the GMM (see Appendix B for details). Lines indicate the contours of Gaussian distributions. (B) Dependence of $\tau$ on $t_b$ for cells classified into subpopulations with longer (green) and shorter (magenta) doubling time. A yellow and black striped bar indicates subjective night, and yellow bar indicates subjective day. (C) Reconstructed histograms of doubling time in experiment [13] obtained with the current method. (D) Histograms of time at division $t_d$ after shifting time at birth and division by 12 hours. Note that subjective day is now between 0 and 12 hours, and subjective night is between 12 and 24 hours.

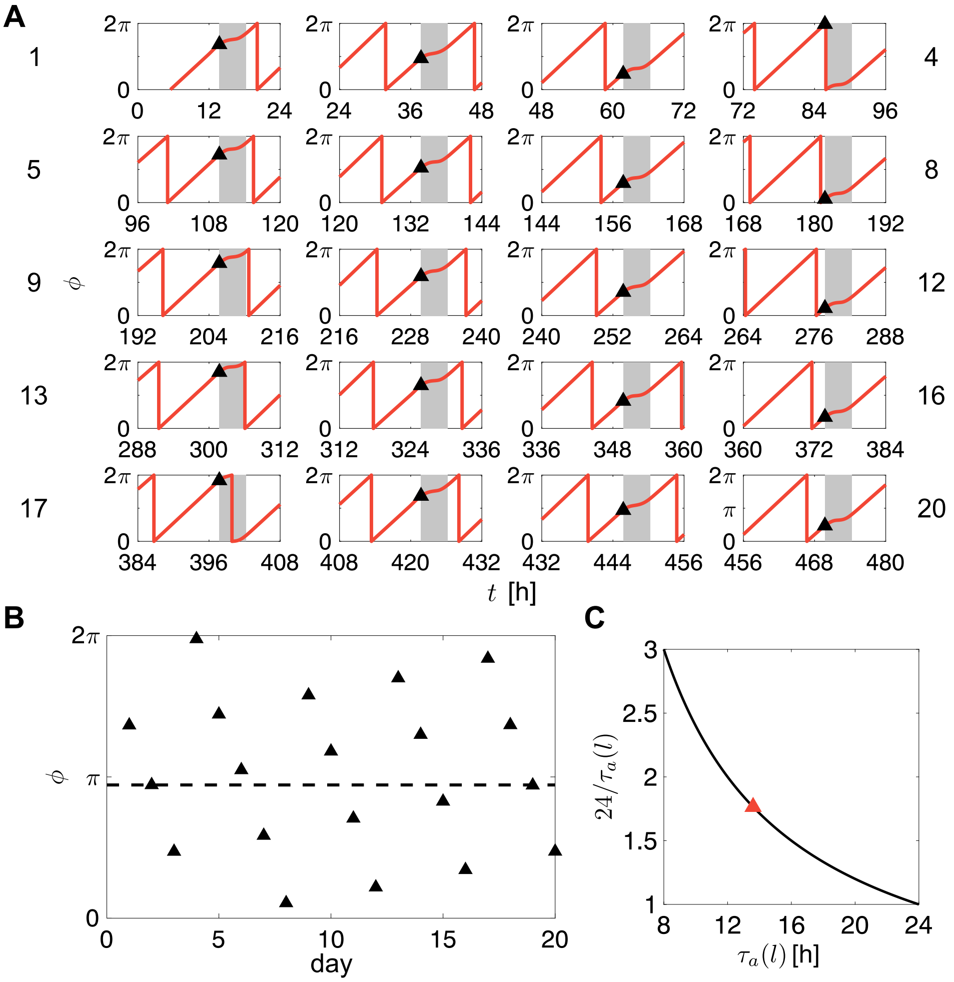


Figure S2: Changes in cell phase $\phi$ at the onset of circadian gating. (A) Time series of cell phase for 20 circadian cycles. Gray shades show circadian gating. Black triangles indicate cell phase at the onset of gating. Numbers next to panels indicate days. Cells are subject to gating mostly once per two generations. (B) Cell phase at the onset of gating indicated by triangles in (A). The dotted line is a visual guide to see the periodicity. 30 cell divisions occur during 17 circadian cycles. (C) The number of cell divisions during a circadian cycle. $\tau_a(l)$ is the doubling time determined by the adder mechanism alone. Red triangle shows the doubling time $\tau_0$ (13.6 hours) in this paper.

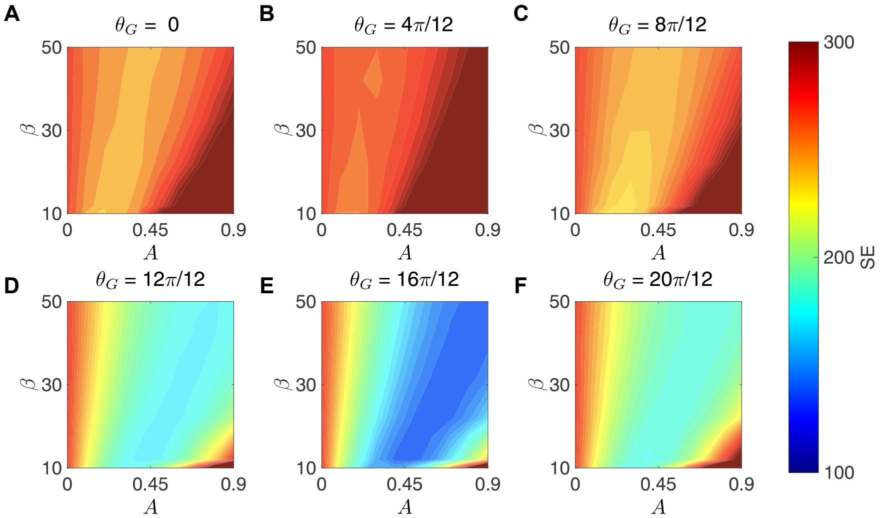


Figure S3: Dependence of square error (SE) in the least-squares fitting on the inhibition strength $A$ and exponent $\beta$ with different values of gating phase $\theta_G$ in the gating function. The color code indicates the value of SE. See the Appendix D for the definition of SE. (A) $\theta_G = 0$, (B) $\theta_G = 4\pi/12$, (C) $\theta_G = 8\pi/12$, (D) $\theta_G = 12\pi/12$, (E) $\theta_G = 16\pi/12$, and (F) $\theta_G = 20\pi/12$. In the main text, we fix $\theta_G = 16\pi/12$ and search $A$ and $\beta$ values that minimize SE.

**Algorithm 1** Calculation of $t_b,\ t_d,\ \tau,\ D,\ l$ with the gating model

System initialization; $\phi = 0, \theta = 0$
Setting parameter values in table 1
$\tau_0 = 13.6,\ \alpha = \ln 2/\tau_0,\ \Delta = 2.85,\ l = 2.85,\ L = l,$
Compute N generations to obtain time series of cell length and doubling time
**for** $i < N$ **do**
    Calculate $\tau_a(l) = (1/\alpha)\ln(1 + \Delta/l)$
    Solve the following ODEs with ode45 in MATLAB until $\phi$ reaches $2\pi$;
    $d\phi(t)/dt = 2\pi\gamma(\theta)/\tau_a(l),\ dL(t)/dt = \alpha L,\ d\theta(t) = \omega$
    Get the array of $\theta$ and $L$
    Initialization of $\phi$ after division; $\phi \leftarrow 0$
    **if** $i \geq 101$ **then**
        Store the arrays of required variables, such as $L$ and $\theta$, for later use.
    **end if**
    Set the initial values of the next generation by using the final values $L_f$ and $\theta_f$ in the array of $L$ and $\theta$ of the current generation;
    $l, L \leftarrow L_f/2$
    $\theta \leftarrow \theta_f$
**end for**

$\theta_i * 24/2\pi$ and $L_i$ of the first entry of the array are the time at birth $t_{b,i}$ and birth length $l_i$ in $i$th generation, respectively. $\theta_i * 24/2\pi$ of the last entry of the array is time at division $t_{d,i}$. Doubling time of $i$th generation is $\tau_i = t_{d,i} - t_{b,i}$. Subtraction of $l_i$ from $L_i$ of the last entry provides added length $D_i$.
