## Supplemental Table for "Cell size homeostasis under the circadian regulation of cell division in cyanobacteria"

Table S1: Statistics of doubling time $\tau$, added length $D$ and birth length $l$ for different cell types in experiment from Martins et al. 2018 [13]

| | | average ($\mu$) | variance ($\sigma^2$) | C.V. ($\sigma/\mu$) |
|---|---|---|---|---|
| $\tau$ [h] | WT | 14.72 | 10.67 | 0.22 |
| | WT shorter $\tau$ | 11.67 | 4.16 | 0.17 |
| | WT longer $\tau$ | 16.32 | 6.65 | 0.16 |
| | $\Delta kaiBC$ | 13.68 | 3.57 | 0.14 |
| $D$ [$\mu m$] | WT | 3.12 | 0.41 | 0.20 |
| | WT shorter $\tau$ | 2.58 | 0.24 | 0.19 |
| | WT longer $\tau$ | 3.41 | 0.26 | 0.15 |
| | $\Delta kaiBC$ | 2.85 | 0.13 | 0.13 |
| $l$ [$\mu m$] | WT | 3.42 | 0.11 | 0.10 |
| | WT shorter $\tau$ | 3.59 | 0.09 | 0.09 |
| | WT longer $\tau$ | 3.34 | 0.09 | 0.09 |
| | $\Delta kaiBC$ | 3.08 | 0.05 | 0.07 |
